# SegmA: Residue Segmentation of cryo-EM density maps

**DOI:** 10.1101/2021.07.25.453685

**Authors:** Mark Rozanov, Haim J. Wolfson

**Affiliations:** Blavatnik School of Computer Science,Tel Aviv University

## Abstract

The cryo-EM resolution revolution enables the development of algorithms for direct de-novo modelling of protein structures from given cryo-EM density maps. Deep Learning tools have been applied to locate structure patterns, such as rotamers, secondary structures and C*α* atoms. We present a deep neural network (nicknamed SegmA) for the residue type segmentation of a cryo-EM density map. The network labels voxels in a cryo-EM map by the residue type (amino acid type or nucleic acid) of the sampled macromolecular structure. It also provides a visual representation of the density map by coloring the different types of voxels by their assigned colors. SegmA’s algorithm is a cascade of CNNs and group rotational equivariant CNNs. A data gathering algorithm was designed for creating datasets that will give best results when used for SegmA’s training. At resolution of 3.2°*A* SegmAs accuracy is 80% for nucleotides. Amino acids which can be seen by eye, such as LEU, ARG and PHE, are detected by SegmA with about 70% accuracy. In addition SegmA detects regions where the exact labeling is of low confidence due to resolution, noise, etc. Removing those “unconfident” regions increases the amino acid detection accuracy to 80% The SegmA open code is available at https://github.com/Mark-Rozanov/SegmA_3A/tree/master.

## 2 Introduction

Knowledge of the three dimensional structure of a protein is a major tool in elucidating its functionality. Cryo-electron microscopy (cryo-EM) (Callaway (2017)) has become a key experimental method for obtaining 3D visualization maps of large molecular structures. The “resolution revolution” (Kühlbrandt (2014)) in cryo-EM has led to an increasing number of near atomic resolution density maps deposited in the EM databank EMDB (Lawson et al. (2016)), making cryo-EM one of the leading experimental methods for determining protein structure. A microscope samples tens of thousands of images which are sorted, aligned and averaged to produce a 3-D density map volume. Afterwards a modelling process is applied to calculate atomic coordinates from this 3D map. The typical modelling process is mainly manual, aided by computational modelling tools.

De-novo modelling methods process a cryo-EM map and obtain an atomic structure when homologous structures are not available. Traditional de-novo cryo-EM modelling algorithms utilize methods like template matching, 3D image analysis and energy minimization. Lasker et al. (2007), Lindert et al. (2012), Baker et al. (2011), Terashi & Kihara (2018), Chen et al. (2016), Liebschner et al. (2019), DiMaio et al. (2011) are examples of such methods. RENNSH (Ma et al. (2012)), Pathwalking (Chen et al. (2016)), SSELEarner (Si et al. (2012)) utilize traditional machine learning (ML) methods in structure determination. Those include among others kNN - k nearest neighbor clustering (RENNSH), SVM - support vector machines (SSE learner) and k-means (Pathwalking).

Recently Deep Learning (DL) tools started to play an important role in cryo-EM modeling. Convolutional Neural Networks (CNN) and their variations perform well on the task of secondary structure detection (Maddhuri Venkata Subramaniya et al. (2019), Moritz et al. (2019)). One of the first attempts to get structural insight at the amino acid level was AAnchor(Rozanov & Wolfson (2018)). AAnchor estimates amino acid centers and types using a 3D CNN. Only residues recovered with high confidence (nicknamed anchors) are reported. *A*^2^-Net (Xu et al. (2019)) is a two-stage modeling algorithm: a CNN for pattern recognition is followed by a Monte Carlo search to determine the protein structure. DeepTracer (Pfab et al. (2021)) uses a cascaded convolutional neural network to predict secondary structures, amino acid types, and backbone atom positions. Structure Generator (Li et al. (2020)) uses a CNN to estimate amino acids with their poses, which are then fed to a graph convolutional network and a recurrent network to obtain the 3D protein structure. For a comprehensive review of Deep Learning methods in cryo-EM modelling we recommend Si et al. (2021) and Li et al. (2020).

Despite recent advances, fully automated protein structure determination from a cryo-EM map is still challenging. There are natural factors that limit the precision of an atomic structure estimation. Among those is the existence of regions of degraded quality in an experimental cryo-EM map caused by molecular flexibility or resolution degradation at the periphery of a molecular complex (Joseph et al. (2020), Si et al. (2021)). Another factor is the variation in residue size and conformation for different amino acid types. As a result residue recognition precision strongly depends on the amino acid type (Rozanov & Wolfson (2018)). The third factor to be mentioned is protein-RNA and protein-DNA complexes. Modelling of such a complex requires separation between amino acids and nucleic acids.

We address the three above mentioned challenges by a segmentation approach. In this segmentation task the output is a map of the same dimension as the input map where each voxel in the output map is coloured according to the amino acid type it belongs to (see Fig. 10).

Namely, the proposed DL algorithm produces a coloured visualization of an input cryo-EM map. In SegmA’s output the residue conformation and boundary are seen in addition to the residue position and type. Voxels belonging to nucleic acids are marked by a different color, which enables separation of RNA/DNA from the proteins and the background. SegmA also assigns a confidence label to each voxel. By selecting only confident voxels for presentation, a user obtains a more precise segmentation at the price of the total amount of voxels that are resolved. Previous studies have highlighted some of the best practices in DL processing of a cryo-EM map:

- Use of cascade structures has advantages for unbalanced data. In this approach the first stage classifier rejects most false negatives and the successive classifiers perform exact labeling.
- Fully Convolutional U-Net architecture is a key to fast processing of a 3D image.
- A reliable data normalization technique is crucial due-to large data distribution variations (Fig. 1).

**Figure 1:**
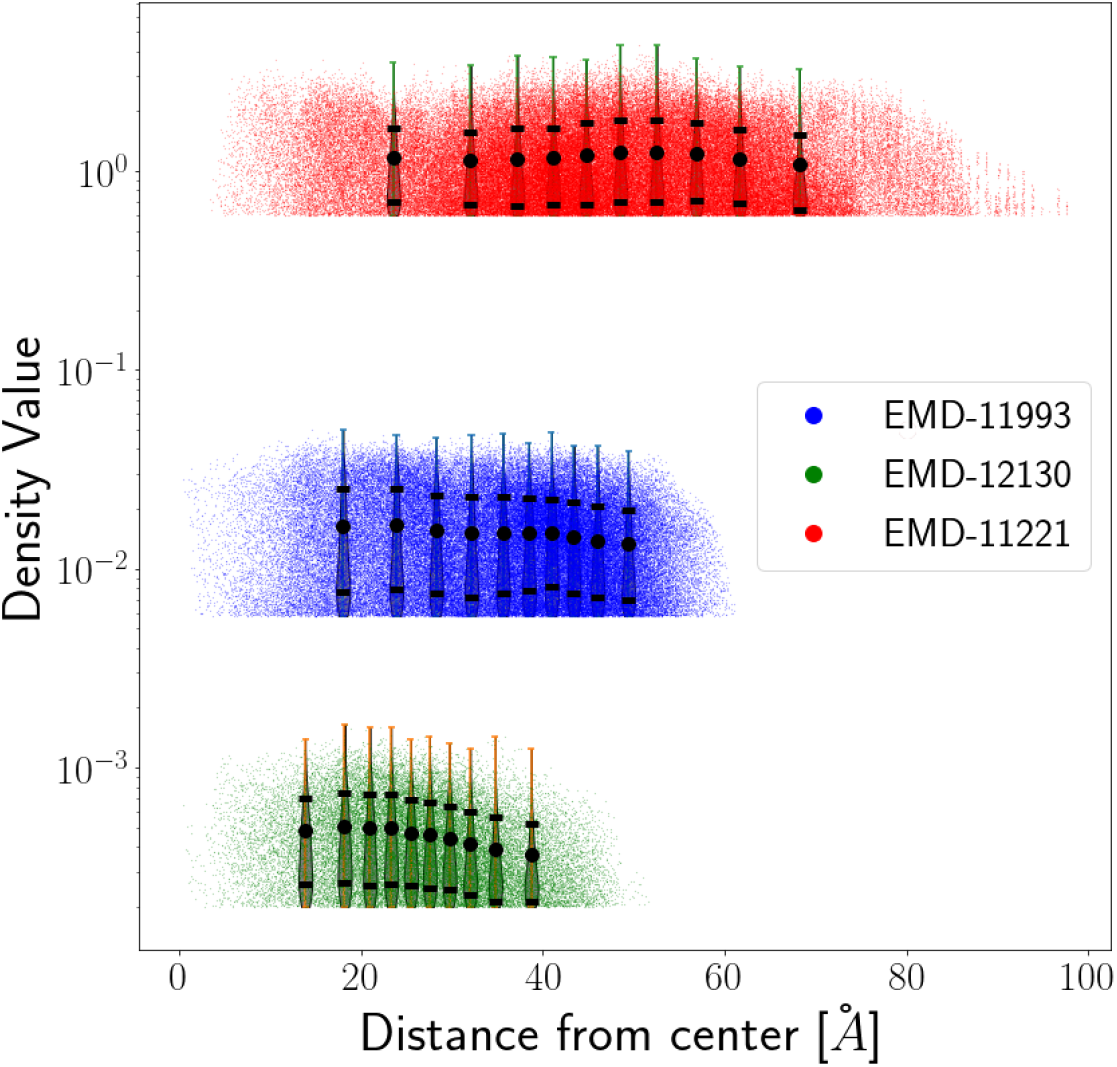
Density value distribution for three cryo-EM maps. Violin plots show the density distribution at a given distance from the map center. Errorbars show mean and standard deviation value. The density distribution significantly differs from one map to another. Moreover density distribution parameters vary as a function of the distance to the map center, while each map has its own pattern.

SegmA incorporates the approaches mentioned above. In addition SegmA incorporates some novel ap-proaches:

- SegmAs segmentation is based on the Group Convolutional Equivariant Network (G-CNN), (Cohen & Welling (2016)), to leverage rotation equivariance of a protein pose in a cryo-EM map.
- SegmA has the ability to estimate the regions where correct prediction is available and report results only in those regions.
- SegmA utilizes a simple and fully automatic input data normalization strategy.

An iterative data gathering algorithm was implemented to create a training data set which results in best segmentation performance.

At resolution of 3.2°*A* SegmAs accuracy is 80% for nucleotides. Visually distinct amino acids, such as LEU, ARG and PHE, are detected by Segma with 70% accuracy and about 80%, if only confident regions are considered. The SegmAs open code is available at https://github.com/Mark-Rozanov/SegmA_3A/tree/master.

## 3 Methods

### 3.1 Map Processing

Given a 3D electron density map of a protein/macromolecular assembly, our task is to assign a label to each voxel within the map. The assigned label should specify that the voxel belongs to one of the following 23 types: one of the 20 amino acids (1 *−* 20), nucleotide (21), background (0), or unconfident (22), where “background” denotes voxels which belong neither to proteins nor to amino acids and “unconfident” denotes voxels which have not been labeled confidently enough to be considered. The algorithm is executed by a cascade of three neural networks - see Figure 2.

**Figure 2:**
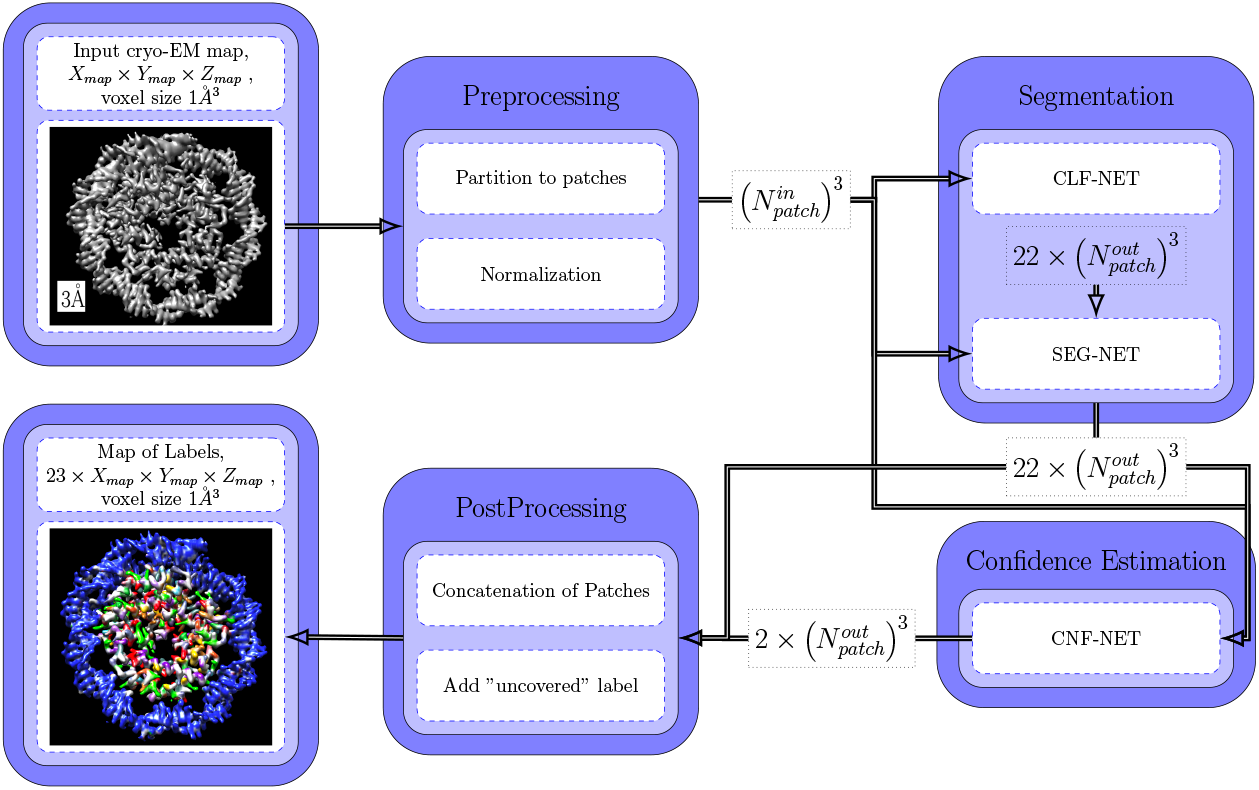
SegmA’s processing scheme. In the **preprocessing** phase an input map is partitioned to overlapping volumetric patches and each patch is normalized. **Classification Net** (CLF-NET) assigns primary labels to each voxel, except for the boundary regions of a patch. **Segmentation Net** (SEG-NET) assigns final labels based on the primary labeling of nearby voxels. **Confidence Net** (CNF-NET) analyses the result of SEG-NET and marks voxels whose labeling is of low confidence. In the **postprocessing** phase all patches are concatenated in a full size map. Voxels marked as “uncorrect” by CNF-NET are reported with the label “unconfident” (22). The rest are reported with the labels assigned by SEG-NET.

The initial labeling of the processed volume is performed by the Classification Net (CLF-NET). The CLF-NET is a group convolutional network, which has the rotation equivariance property (Cohen &Welling (2016)), i.e., its features are invariant to rotation of the input cryo-EM map. The Classification Net receives as input a normalized volumetric patch of the cryo-EM map and outputs the labels of each voxel within this patch. CLF-NET has a receptive field of 10°*A*^3^.

The results of CLF-NET are fed to the Segmentation Net (SEG-NET), which performs the final labeling of the voxels. SEG-NET is a U-net CNN (Ronneberger et al. (2015)) which combines the cryo-EM values of the input voxels with the results of CLF-NET on the neighboring voxels. The Segmentation Net has two inputs: a normalized patch of the cryo-EM map and the result of CLF-NET for this patch. The output is the final label of each voxel, which takes into account the neighborhood information. SEG-NET has a receptive field of 15°*A*^3^.

Finally, the Confidence Net (CNF-NET) evaluates the results of SEG-NET. It assigns a binary label (1 - correct, 0 incorrect) to each voxel. Only “correct” voxels are reported, the rest are marked with label 22 (unconfident). The Confidence Net is a rotation equivariant convolutional network. It has a receptive field of 15°*A*^3^ and 24 channels of input (the source cryo-EM map and 23 output channels of the SEG-NET).

### 3.2 Input Normalisation

Experimental cryo-EM maps suffer from variation in their density value distributions. While voxels with higher density should mark the locations of atoms, density values of similar molecules may differ significantly from one map to another (Lawson et al. (2020)). This is due to lack of standardization combined with multiple factors affecting the density values, such as the microscope type and reconstruction software. Online databases such as EMDataResource (Lawson et al. (2016)) assign different threshold values for each map for the interpretation of atom locations. Fig. 1 shows density values for three test maps on a logarithmic scale. Each of the three shown cases has a unique density distribution which is different from one map to another. Moreover the density distribution varies as a function of the distance of a voxel to the map center. This variation does not have any recognisable pattern.

Machine learning algorithms usually apply a data normalisation procedure which ensures that the training data and the test data are of similar distribution. Cascaded-CNN (Moritz et al. (2019)) has a five stage normalisation algorithm based on ordered statistics and manually selected thresholds. Emap2Sec (Maddhuri Venkata Subramaniya et al. (2019)) normalises each voxel in the map to the range 0 *−* 1 by a linear transformation. AAnchor (Rozanov & Wolfson (2018)) adjusts the mean and standard deviation of each sample box of size 10°*A*^3^. Normalising each sample box prevents using a fully convolutional network on the whole map and therefore is time consuming. Having said that, note that normalising a whole map is not efficient, since the density distribution changes within a map. Moreover, a typical map has a large “background” volume which does not contain any atoms. SegmA applies a trade-off approach. A map is divided into overlapping patches of size 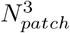, where each patch is normalised to mean value of 0.5 and standard deviation of 0.15. A fully convolutional neural network is applied separately on each patch. Large values of *N*_*patch*_ result in fast running time at the cost of density distribution variability. Based on empirical estimations, SegmA uses *N*_*patch*_ = 45°*A*.

### 3.3 Group CNN for the Amino Acids Classification

A function *f* (*x*) is said to be **equivariant** to a function *g* if *f* (*g*(*x*)) = *g* (*f* (*x*)). A function *f* (*x*) is said to be **invariant** to a function *g* if *f* (*g*(*x*)) = *f* (*x*). Note that the matrix convolution operation is equivariant to translation: a shift in the input leads to a corresponding shift in the output. Due to the translation equivariance, conventional CNNs classify structural patterns regardless of their location without additional effort.

However many classification tasks, especially in 3D, should also accommodate the *rotational symmetry* property: a rotation of an object should not change its classification. Moreover, sometimes rotational symmetry holds for local volume fragments, for example, a rotation of an amino acid by 90^°^ should not alter its classification. The most popular approach to deal with angular rotations is an augmentation of the training dataset with rotated versions of the input. In this case a CNN “duplicates” learning features for “all” possible rotations. The obvious disadvantages of such an approach are more critical in 3D, due to the large input data size and the 3D space of rotations.

Group convolutional neural networks (G-CNNs), introduced in Cohen &Welling (2016), have an architecture which exploits the rotational symmetry. A layer in a G-CNN performs operations which are equivariant to rotations, i.e. 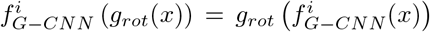, where *g*_*rot*_ is a rotation function and 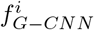 is the *i*-th layer of the G-CNN network. A 3D classification G-CNN processes a 3D input matrix and outputs a scalar label, therefore G-CNN is **rotation invariant**: *f*_*G-CNN*_ (*g*_*rot*_ ((*x*)) = *f*_*G-CNN*_ (*x*) where *fG-CNN*: *R*^*N*×*N*×*N*^ → {1, …*N*_*labeles*_} is a G-CNN classification function.

A G-CNN layer is built on the basis of a symmetry group of a discrete set of rotations. Conventionally, a discrete set of rotations by 90^°^ in each of the 3 dimensions is used, because those are actually permutations of 3D matrix elements and no interpolation and matrix multiplication are required. Table 1 lists the rotation groups used in this work.

**Table 1:** 3D Rotation Groups.

|  |  |  |
| --- | --- | --- |
| Klein | 4 Rotations | Set of all $180^\circ$ rotations around $x, y, z$ axes. |
| T4 | 12 Rotations | Tetrahedral group. Set of all two-axes rotations by $90^\circ$. |
| S4 | 24 Rotations | Set of all $90^\circ$ rotations around $x, y, z$ axes. |

From the implementation point of view a group convolution layer outputs *N*_*rot*_ 4D matrices of dimension *N*_*chan out*_ *× L × W × H*, where *N*_*chan out*_ is the number of output channels and *L, W, H* are the length, width and height of a filter. Cohen &Welling (2016) provide extended explanation of group CNNs, including the theoretical basis and implementation issues. For understanding group CNNs in three dimension we recommend Worrall & Brostow (2018) and Winkels & Cohen (2019). SegmAs G-CNN implementation was based on the open source code of Romero et al. (2020) and Worrall & Brostow (2018).

The architecture of the Classification Net is presented in Fig. 3a. CLF-NET assigns a label to the center of an 11°*A*^3^ cube. CLF-NET has 22 label types: 20 amino acids, nucleotide and background.

**Figure 3:**
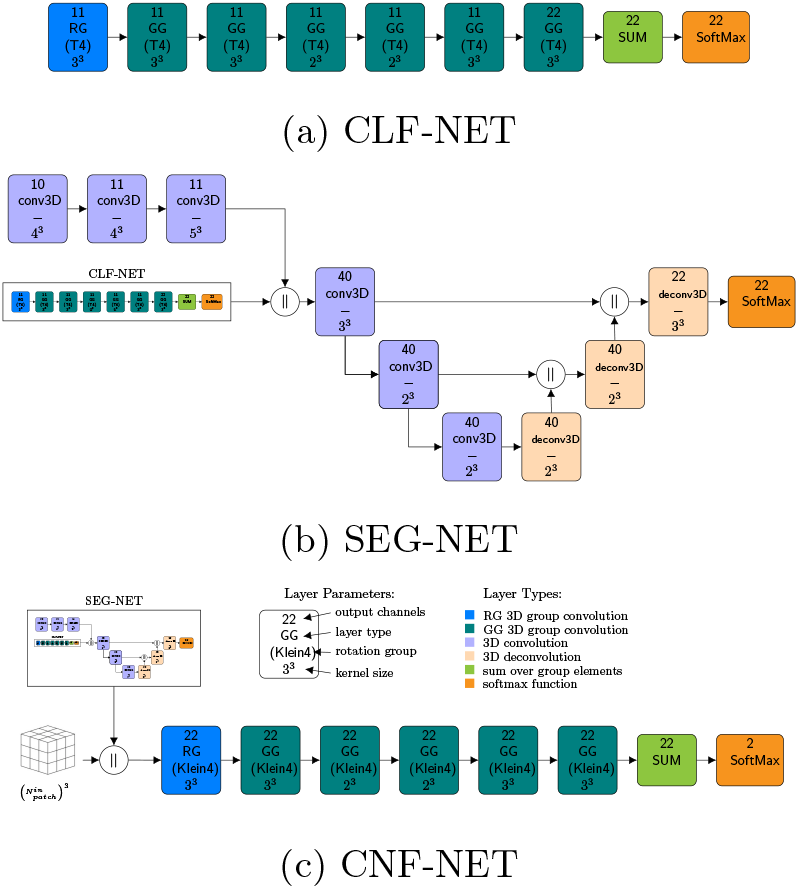
(a) The Classification Net (CLF-NET) architecture based on the T4 group. This is a typical group CNN network. A GG convolution layer performs convolution on a group of *N*_*rot*_ rotations. The very first layer is a RG convolution which inputs a 3D map and outputs a group. The sequence of GG layers is followed by a summation layer. (b) The Segmentation Net (SEG-NET) architecture based on the U-net approach. (c) The Confidence Net: an equivariant network based on the Klein group.

The architecture of the Segmentation Net (SEG-NET) is presented in Fig. 3b. SEG-NET utilizes the U-net network architecture. First proposed for the analysis of 2D biomedical images by Ronneberger et al. (2015), nowadays U-net is one of the most powerful architectures in image segmentation tasks, both in 2D and 3D (Siddique et al. (2020)). Typical U-net architecture consists of a contraction path and an expansion path. The contraction path, also known as the encoder resembles a convolution network and provides classification information. SEG-NET has two contraction paths: one is a conventional sequence of convolutional layers, and the other is the pretrained CLF-NET. The outputs of the two paths are concatenated and fed into the last layer of the encoder. The expansion path or the decoder consists of a sequence of deconvolution (or inverse convolution) layers. An input to a deconvolution is the concatenation of output from the previous layer with the output from one of the convolution layers from the encoder.

The Confidence Network (CNF-NET) is a typical group CNN and its architecture is like that of CLF-NET. The input is a 3D map with 23 channels, which is the normalized patch of a cryo-EM map concatenated with the result of SEG-NET (22 pseudo-probabilities). The output is a binary label which asserts the SEG-NET labeling: 1 - correct, 0 incorrect.

All networks use the rectified linear (ReLU) activation function. The networks are implemented as Fully Convolutional Networks (FCNs, Long et al. (2015)), so they run on a variable input size. The FCN architecture reduces the running time of SegmaA: training and testing are done on a large map patch and no sampling or sliding window is required.

We first train the rotation equivariant CLF-NET, afterwards SEG-NET training starts from the pretrained values of CLF-NET. The CNF-NET is trained last on the output of SEG-NET.

### 3.4 Data Processing

The **preprocessing** of the training data consists of cropping, augmentation, interpolation, normalization and labeling. Test data preprocessing consists of interpolation and normalization. Given a cryo-EM map and a fitted atomic model, the cropping procedure crops out the cryo-EM map edges which are far from the fitted model. No data augmentation was needed, since rotation invariance is achieved by using group equivariant CNNs. The interpolation phase modifies the voxels size of a map to 1°*A*^3^. The labeling procedure assigns to each voxel one of the 22 labels (see Fig. 4 for details).

**Figure 4:**
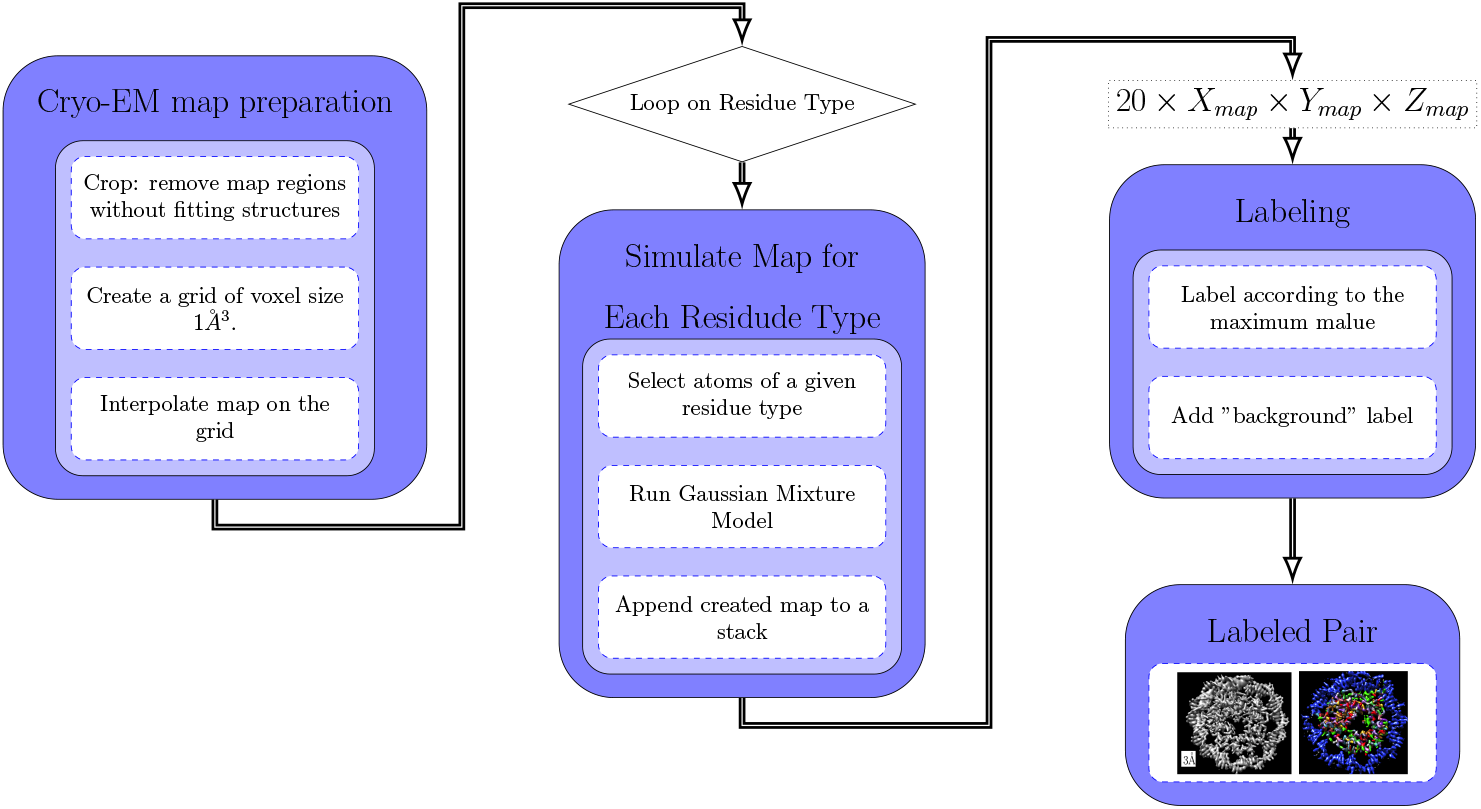
Labeling Procedure: First, the map is cropped and interpolated to voxel size of 1°*A*^3^. Afterwards the pdb2mrc routine (Tang et al. (2007)) is applied only for atoms of a specified residue. pdb2mrc creates a 3D density map which is Gaussian Mixture, where the Gaussian centers are the atom centers and amplitudes are proportional to the atomic number. A residue type whose GMM map has a maximum value at an (*i, j, k*) voxel is the voxel label. Voxels with a maximal GMM value below a threshold are labeled as background (0).

An amino acid type (or nucleotide) is assigned to a voxel which is in the proximity of an atom of the residue of this type. A voxel with no residue atom in its proximity gets the label 0 - background.

In the final stage the SEG-NET labeling of the map is modified according to the CNF-NET output. Namely, voxels marked by CNF-NET as “correct” are reported with the original SEG-NET labeling, the rest receive the label “unconfident”.

### 3.5 Database creation

Despite the fact that there are thousands of maps with fitted structures available online (Fig. 5a), they can’t be used in the training without filtering. This is because some of them contain parts of imprecise fit, missing atomic structure or missing map density (Fig. 5b), as those maps were created for understanding biological functionality and not as entries of a ML database.

**Figure 5:**
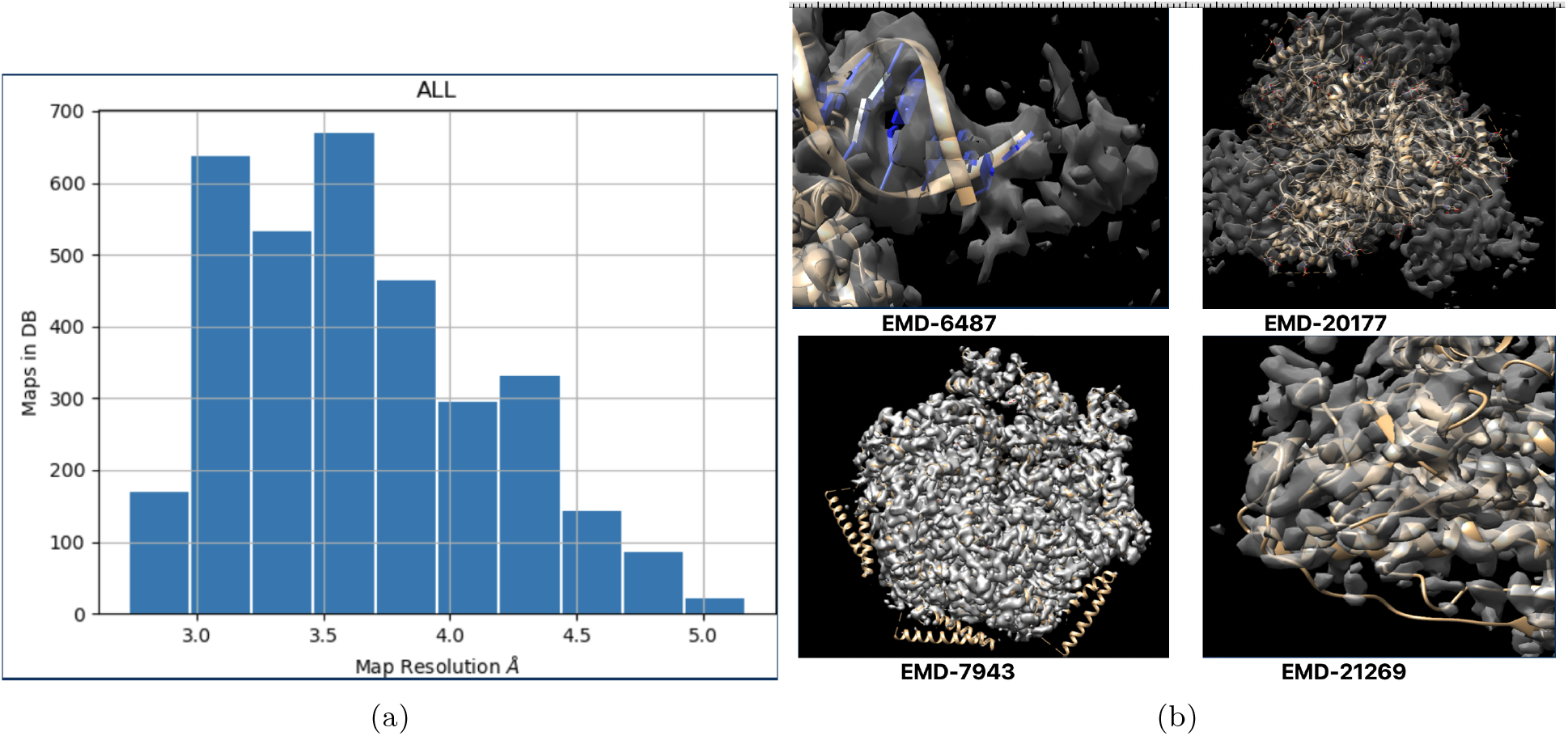
(a) Reported resolution distribution of single particle cryo-EM maps with fitted structures as downloaded from EMDB at time of creation. (b) Examples of imperfect fitting: EMD-6487 (Ru et al. (2015)) - some parts of the DNA helix are missing, EMD-20177 (Escolano et al. (2019)) - map with unfitted parts, EMD-7943 (Yu et al. (2018)) - cryo-EM map does not covers helices, EMD-21269 (Yang et al. (2020)) - the fitting of some loops is not precise.

Existing Deep Learning methods utilized different approaches for gathering maps in a training dataset: AAnchor (Rozanov & Wolfson (2018)) and *A*^2^-Net (Xu et al. (2019)) manually selected maps for the training size. C-CNN (Moritz et al. (2019)), Emap2Sec (Maddhuri Venkata Subramaniya et al. (2019)) and Rotamer

Net (Li et al. (2020)) were trained on simulated maps. DeepTracer (Pfab et al. (2021)) was trained on 1800 maps without selection. SegmA’s training dataset was gathered using an iterative procedure inspired by the well-known co-training and tri-training methods (Zhou & Li (2005), Zhu (2008), Saito et al. (2017)). The purpose of the procedure is to filter out data samples which may cause the training to go in the wrong direction. A data sample is a normalized patch of a cryo-EM map, see (Sec. 3.2) for details. The iterative data gathering procedure is presented at Alg. 1. We start from initially trained CLF-NET and two datasets: the *candidates* set which is all the patches from all maps in a given resolution span and the *validation* set which consists of a small amount manually examined maps at target resolution. The algorithm aims to extract an appropriate patches from candidates set, such that CLF-NET when trained on those patches, will have best performance on the validation set. At each iteration a new training dataset is created from the candidate samples which have a loss lower then LP percentile (typical 70% or 75%) of the loss of the validation set. Then the CLF-NET is retrained on the new dataset and next iteration starts. As the training dataset improves, CLF-NET predictions are more accurate and the procedure converges - Fig. 6.

**Figure 6:**
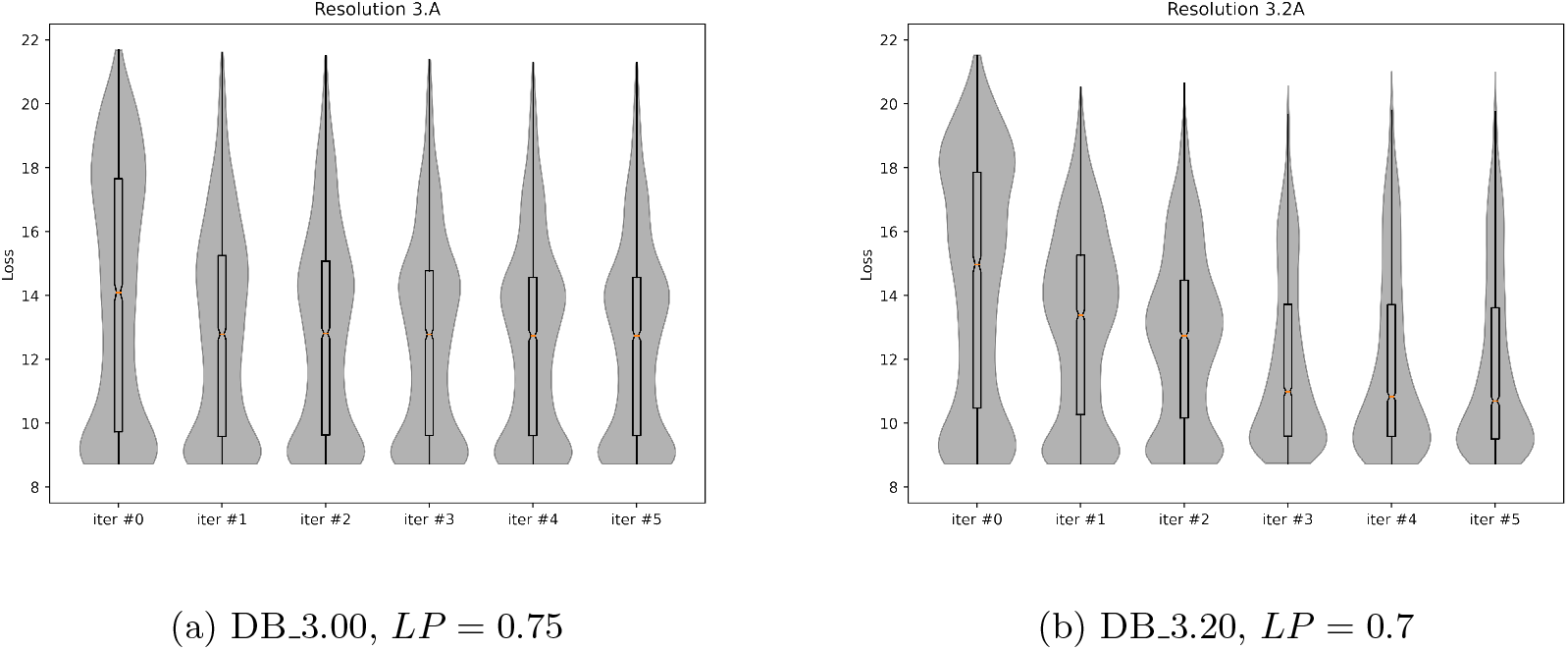
Loss Distribution of Validating set at the start of each iteration of Alg. 1. The boxes show 25% and 75% percentiles and the red line shows the median value.

## 4 Results and Discussion

### 4.1 Selecting an Optimal Rotation Group

A small scale experiment was performed to determine the rotation group to be used for CLF-NET. The Classification Net was trained and tested in 4 different configurations. Three group convolution networks for each of the symmetry groups (Klein, T4 and S4) as well as for the AAnchor configuration without rotation equvariance were tested on a small dataset of 15 training maps and 3 test maps in resolution range 2.9°*A −* 3.1°*A*. All four configurations had a similar amount of learnable parameters. Table 2 summarizes the performance of CLF-NET and SEG-NET for the four different configurations of CLF-NET.

**Table 2:**
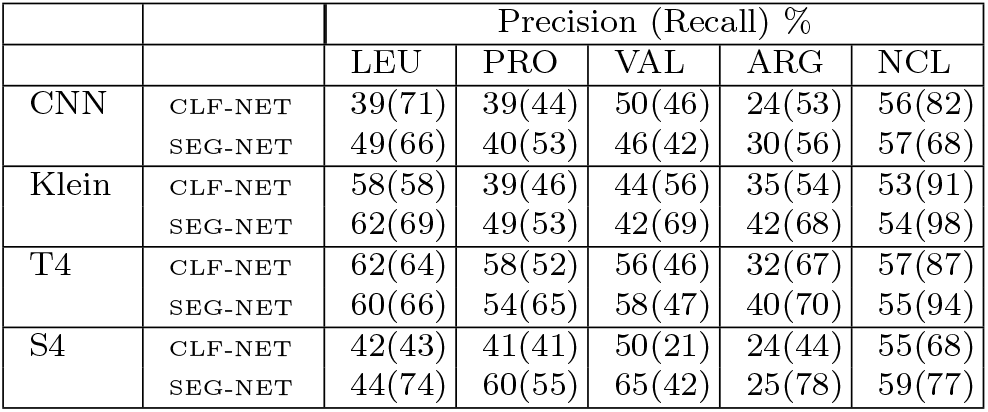
CLF-NET performance for various 3D groups. 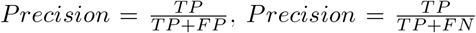. *TP* is a number of true positives. *FP* is a number of false positives. *FN* is a number of false negatives. CNN denotes convolution neural network without rotations. Klein - the group of four 180^°^ rotations. S4 is the group of all possible 90^°^ rotations, 24 members. T4 is the tetrahedral group which consists of 12 two axis rotation by 90^°^

|  |  | Precision (Recall) % |  |  |  |  |
| --- | --- | --- | --- | --- | --- | --- |
|  |  | LEU | PRO | VAL | ARG | NCL |
| CNN | CLF-NET | 39(71) | 39(44) | 50(46) | 24(53) | 56(82) |
|  | SEG-NET | 49(66) | 40(53) | 46(42) | 30(56) | 57(68) |
| Klein | CLF-NET | 58(58) | 39(46) | 44(56) | 35(54) | 53(91) |
|  | SEG-NET | 62(69) | 49(53) | 42(69) | 42(68) | 54(98) |
| T4 | CLF-NET | 62(64) | 58(52) | 56(46) | 32(67) | 57(87) |
|  | SEG-NET | 60(66) | 54(65) | 58(47) | 40(70) | 55(94) |
| S4 | CLF-NET | 42(43) | 41(41) | 50(21) | 24(44) | 55(68) |
|  | SEG-NET | 44(74) | 60(55) | 65(42) | 25(78) | 59(77) |

#### Algorithm 1

Iterative Dataset Gathering. The training set *X*_*t*_ consists of the patches from the candidate set, *X*_*c*_, with obtained classification loss below a threshold. The threshold is *LP* percentile of losses of a validation set, *X*_*v*_.

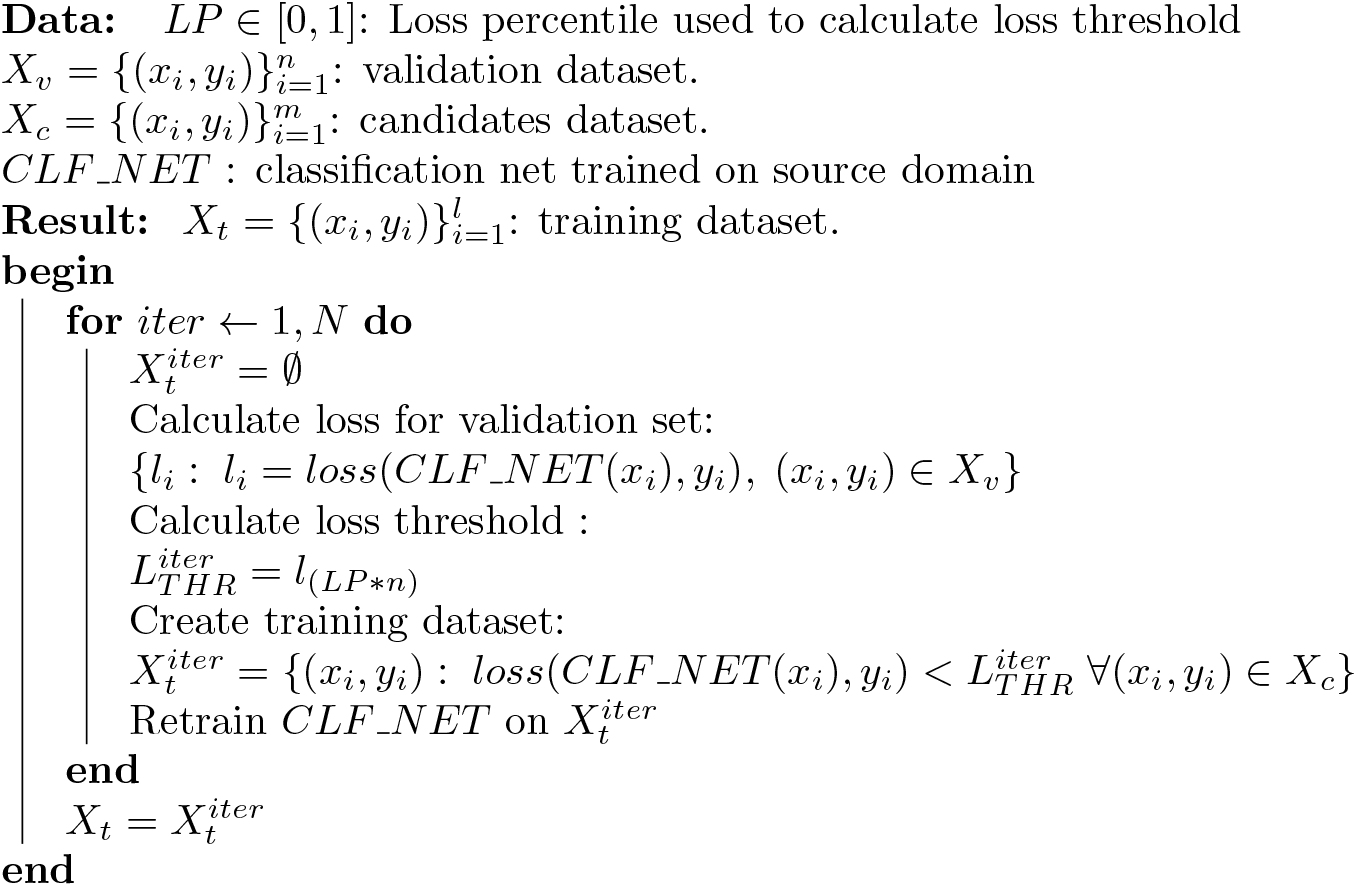

Precision and recall values were calculated for four amino acid types and nucleic acids. One can see that the performance of group convolutional networks is better then performance of the regular CNN network. The performance degradation of the S4 group may be explained by training difficulties. In this configuration each layer has 576 matrices: 24 sets of 24 rotated matrices. The contribution of SEG-NET is notable mostly in the recall values, i.e., more voxels of given amino acid types are detected. The rest of this section shows the results for the T4 group used in CLF-NET. For the CNF-NET Klein group was used due to GPU memory limitations.

### 4.2 The Dataset Gathering

Using Alg. 1 two training datasets were created: DB 3.00 is the dataset dedicated to segmentation of maps with resolutions 2.9 *−* 3.1°*A* and DB 3.20 which is the dataset dedicated to segmentation of maps with resolutions 3.11 *−* 3.33°*A*. See Fig. 7 (c) for the difference between two datasets. Candidate sets were created by partitioning to patches all maps in EMDB with the reported resolution 2.9°*A −* 3.3°*A* for DB 3.00 and 3.0°*A −* 3.4°*A* for DB 3.20. Validation sets were created from a small number of manually chosen maps at resolution 3.00°*A* for DB 3.00 and 3.00°*A* for DB 3.20. For DB 3.00 the initial CLF-NET was trained on ten manually selected maps with reported resolution 2.9°*A −* 3.3°*A*. For DB 3.20 the initial CLF-NET is the CLF-NET from the last iteration of Alg. 1 for DB 3.20. Another difference between the two datasets is the percentile value, *LP*, which was 75% for DB 3.00 and 70% for DB 3.20.

**Figure 7:**
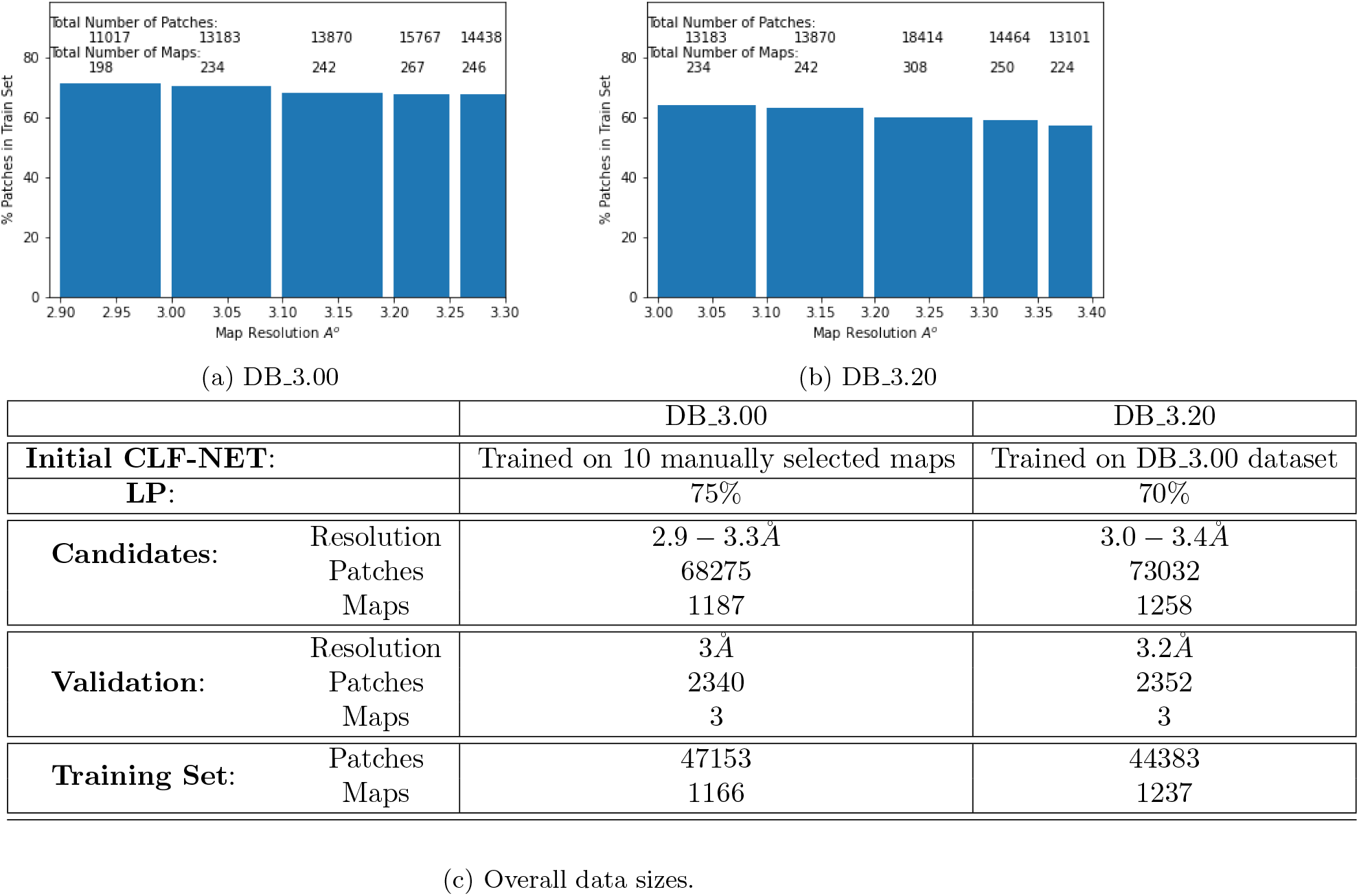
Dataset statistics of DB 3.00 and DB 3.20. (a) (b) show the percentage taken for the training data set per resolution. For both datasets the percentage of filtered out patches maps grows. (c) Datasets sizes before (Candidates) and after (Training Set) applying Alg. 1.

The convergence of Alg. 1 is shown on Fig 6. Surprisingly, DB 3.20 resulted in better results for the validation set. Possible explanation for that is *LP* = 70% used for DB 3.20, as can be seen in Fig 6 a large amount of patches have their loss in region the 75% percentile. Another possible explanation for that is a wider resolution span used for candidates. i.e., DB 3.20 has a large (*≈*500) amount of maps in resolutions 3.3 − 3.4°*A*. A least half of the patches from those map where selected as “good” and inserted in the training set. Fig. 7 shows the obtained dataset sizes and share of the “good” patches per resolution. As expected, the percentage of filtered out patches is higher for the maps with lower resolution.

Since DB 3.20 dataset achieved best performance for both 3°*A* and 3.2°*A* validation maps, the rest of the paper results are for the DB 3.20 dataset.

#### 4.2.1 SegmAs performance evaluation

SegmA’s labeling depends on a number of parameters: local map resolution, amino acid type, density distribution within a map, etc. The performance was evaluated on seventeen test maps of resolutions 2.9 *−* 3.5°*A*.

Usually in a typical experimental cryo-EM map a molecule occupies no more then 10% of total volume, the rest is irrelevant background noise. Including background voxels in the accuracy computation will lead to an unwanted imbalance. In the training dataset those voxels were excluded by cropping a map to a region close to the fitted atomic structures. For the test case, where the atomic fitted structure is assumed to be unknown, background voxels were filtered using the **Recommended Contour Level**, RCL. RCL is a threshold value provided alongside with a cryo-EM map in an online database, Lawson et al. (2016). Voxels with density below this value are assumed to be in the background and are usually not shown to the user. During this paper only voxels with densities above RCL are considered for the performance evaluation.

Fig. 8 shows the performance of SegmA’s three components for Nucleic Acids and three amino acids: Leucine, Arginine and Phenylalanin. Note the contribution of the Segmentation Net to the labeling performance, which is especially significant for large amino acids, like Arginine.

**Figure 8:**
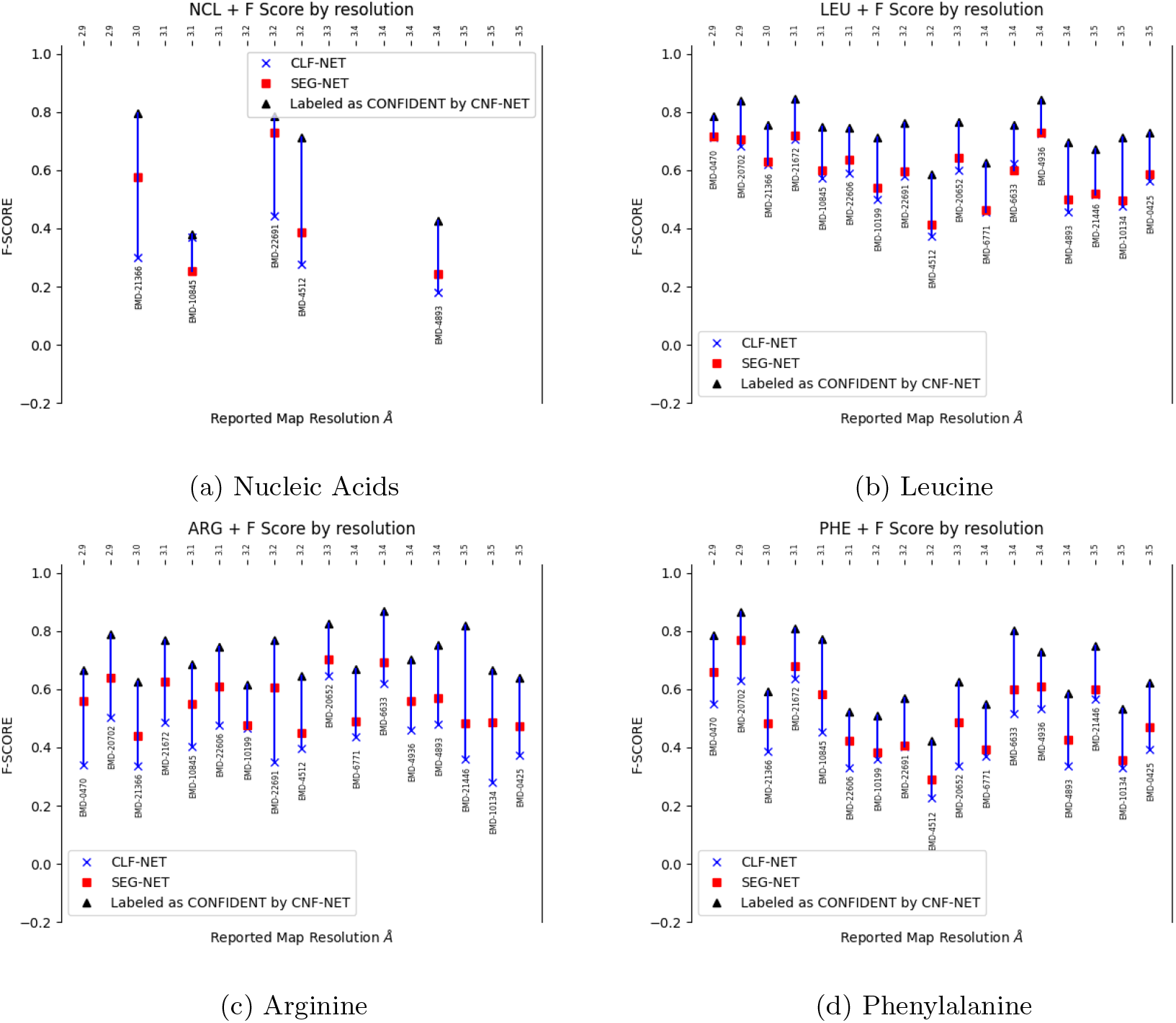
F-Score for sevennteen test maps for Nucleic Acids (a), Leuceine (b), Arginine (c) and Phenylanine (d). 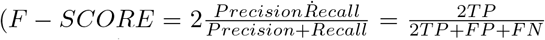. For each map the performance of CLF-NET (blue x’), SEG-NET (red squares) are shown. Black triangles show the improvement in F-Score when only voxels marked as confident by CNF-NET are considered.

For nine test maps at resolutions 2.9 *−* 3.2°*A* we calculated the Confusion Matrix of SEG-NET - Fig. 9. The performance varies for different amino acid types. SegmA achieves around 70% precision for LEU, ARG, ALA, VAL, TYR and above 95% for nucleotides. CNF-NET significantly improves the performance by filtering out false labeled voxels. Fig. 9 shows the rate of filtered voxels by their amino acid type. CNF-NET filters out voxels of amino acids which are hard to detect. As for the amino acid types with high detection rate, and a large amount of false labeled voxels is marked as “unconfident”.

**Figure 9:**
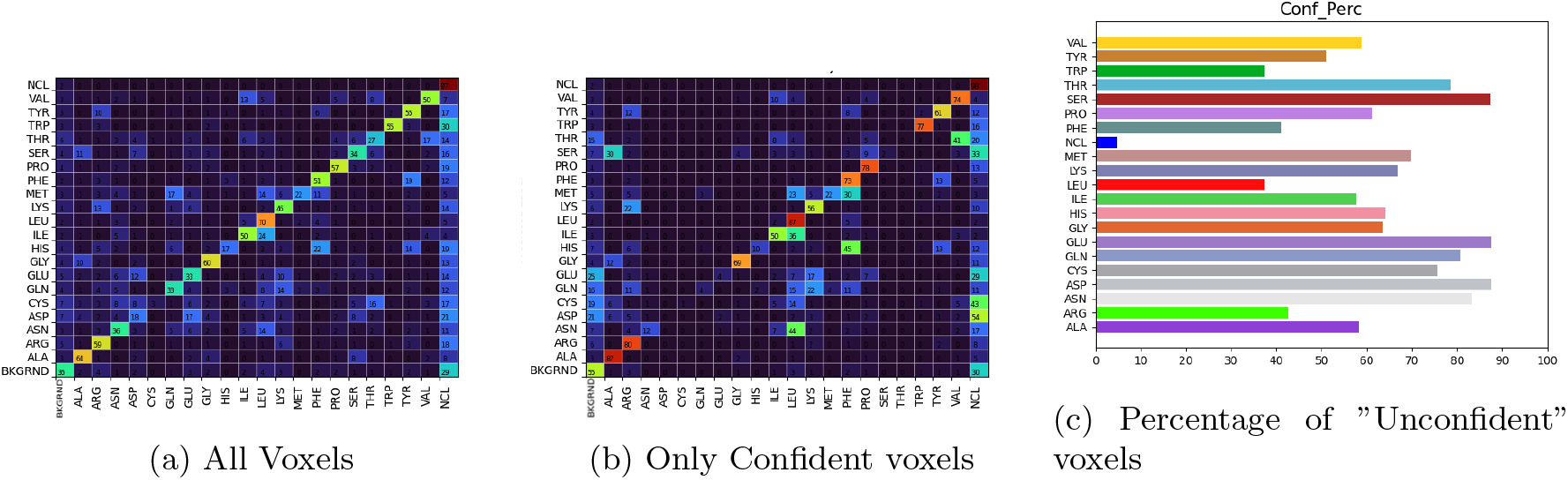
SegmAs performance for nine test maps at resolutions 2.9 *−*3.2°*A*. (a), (b) Row Normalized Confusion Matrix. A row represents a true labeling of a voxel, whereas a column represents SegmAs prediction. (a) shows the output of SEG-NET, (b) shows the results after eliminating unconfident voxels. For example: SEG-NET detected 70% of LEU voxels. Among confident voxels, the rate of detected LEU voxels increases to 87%. (c) Rate of voxels marked as unconfident by CNF-NET. CNF-NET “excluded” almost all voxels belonging to the amino acids which are hard to detect. It also improved the precision for LEU, ARG, ALA, VAL, TYR by at least 15%.

It has been known from years of human eye examination of electron density maps that some amino acids can be misinterpreted. For example Branden & Tooze (2012) mention some “indistinguishable” pairs: THR-VAL, ASP-ASN, GLU-GLN, HIS-PHE and others. SegmA experience on cryo-EM maps is slightly different: most of the mislabeling occurs between amino acids of similar physico-chemical properties. Only LEU-ILE, HIS-PHE and THR-VAL have a considerably high non-diagonal values in the confusion matrix.

SegmAs segmentation results can be best viewed in a 3D visualization tool, such as Chimera or Pymol. Figures 10,11,12,13, show the contour plots generated by Chimera, where each voxel colored according to its amino acid label. Voxels labeled as background are colored black, so are the unconfident voxels on the image showing confident only voxels. Such coloring enables to see an amino acid position and pose. There as some case of amino acids voxels false labeled as nucleotides. This happens mostly on the boundary between the molecule and the background, were local resolution lower. Nevertheless, voxels mislabeled as nucleotides are mostly filtered out by CNF-NET.

**Figure 10:**
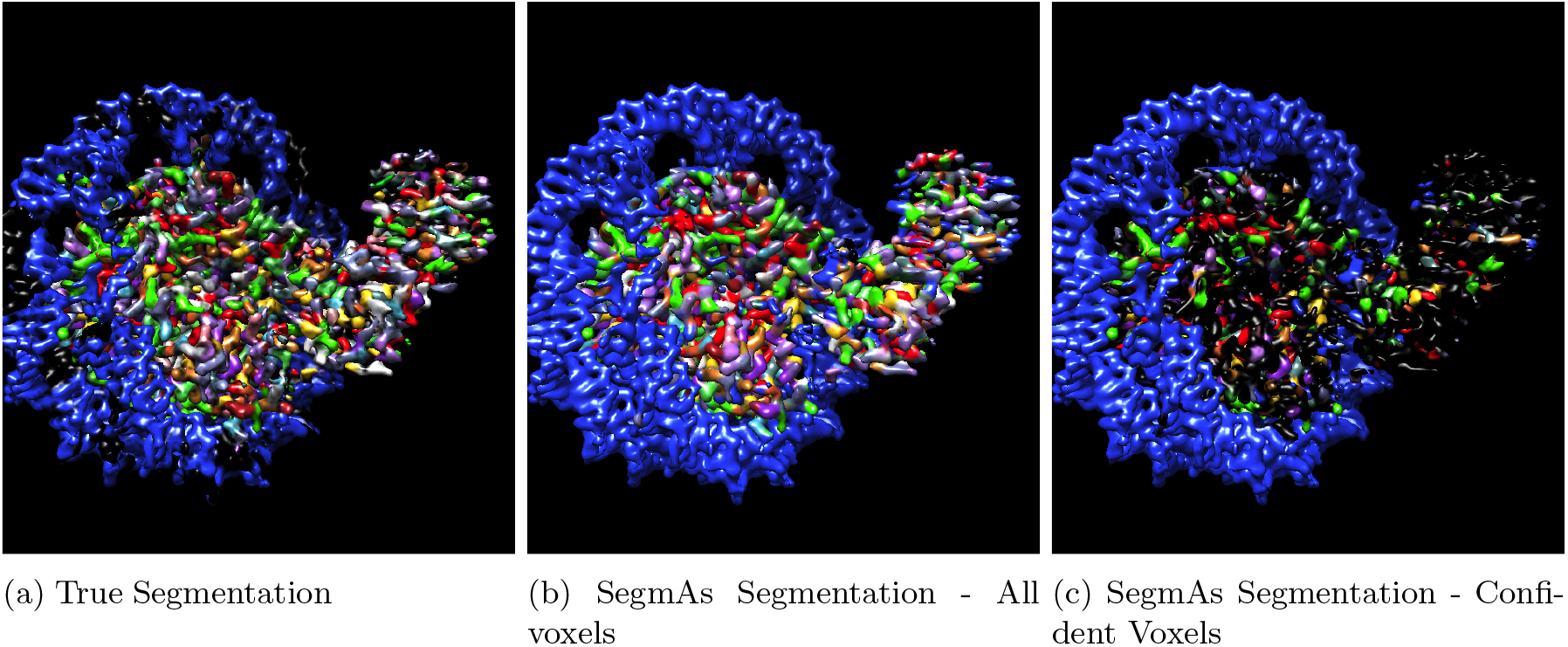
Segmentation of a Nucleosome at 3.2°*A* resolution, EMD-22691. The DNA ring is identified correctly. Most of the mislabeled voxel lie between an amino acids or in a peripheral regions. CNF-NET successfully identifies false labeled voxels.

**Figure 11:**
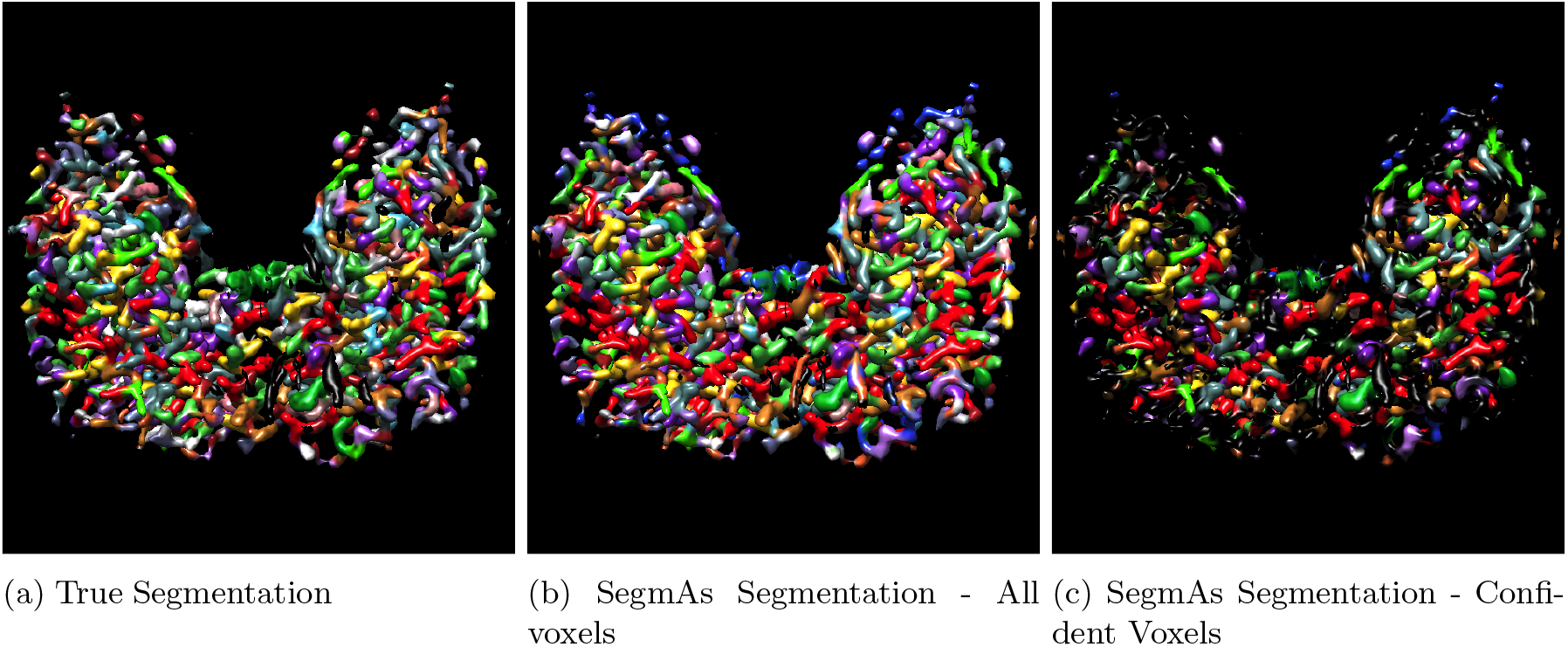
EMD-0470, reported map resolution - 2.9°*A*. Voxels false labeled as nucleotides (blue) are filtered out by CNF-NET.

**Figure 12:**
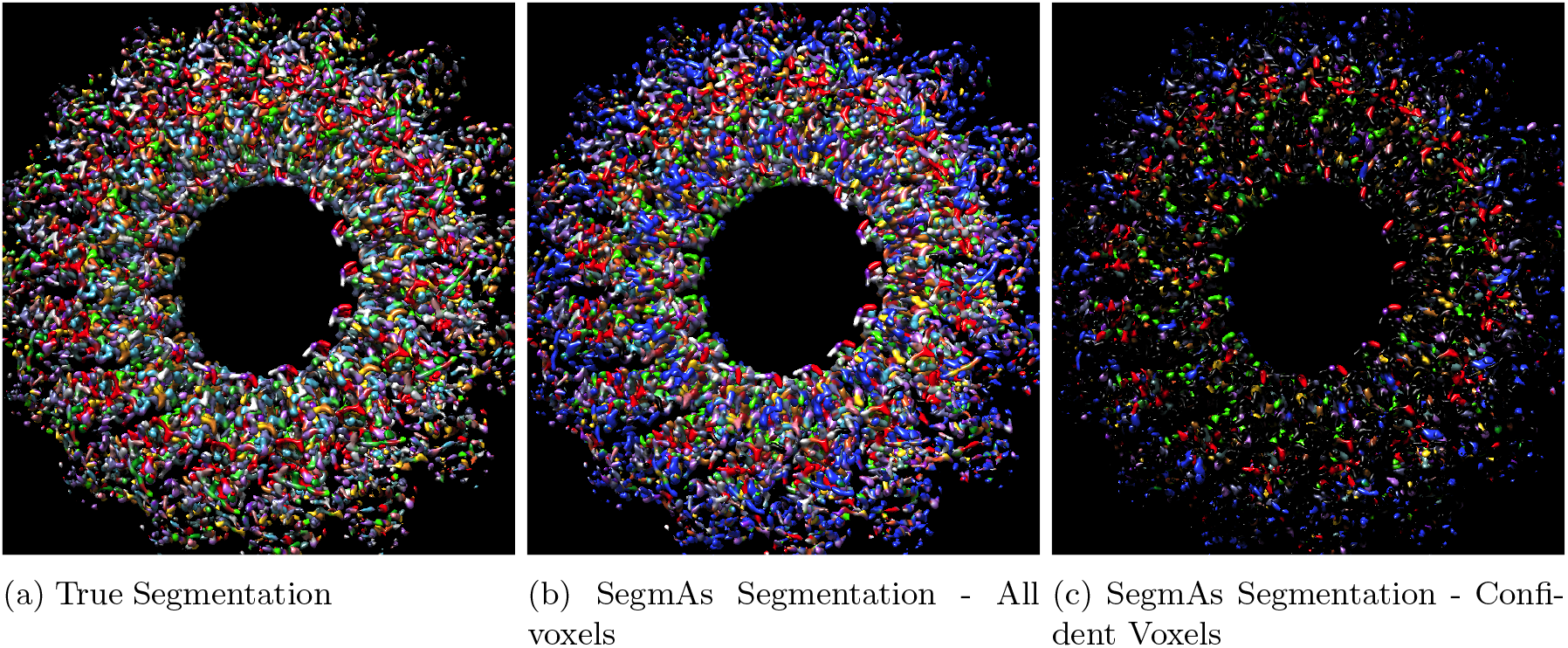
Segmentation of a 3.5°*A* resolution map. EMD-10134, murine perforin-2 ectodomain. While voxels in the periphery were mislabeled as nucleic acids, a majority of them are filtered out by CNF NET. On the inner ring surface SegmA correctly identifies (with confidence) Leucine (red), Isoleucine (green) and Alanine(purple). Unconfident voxels lie mostly in the boundary between amino acids.

**Figure 13:**
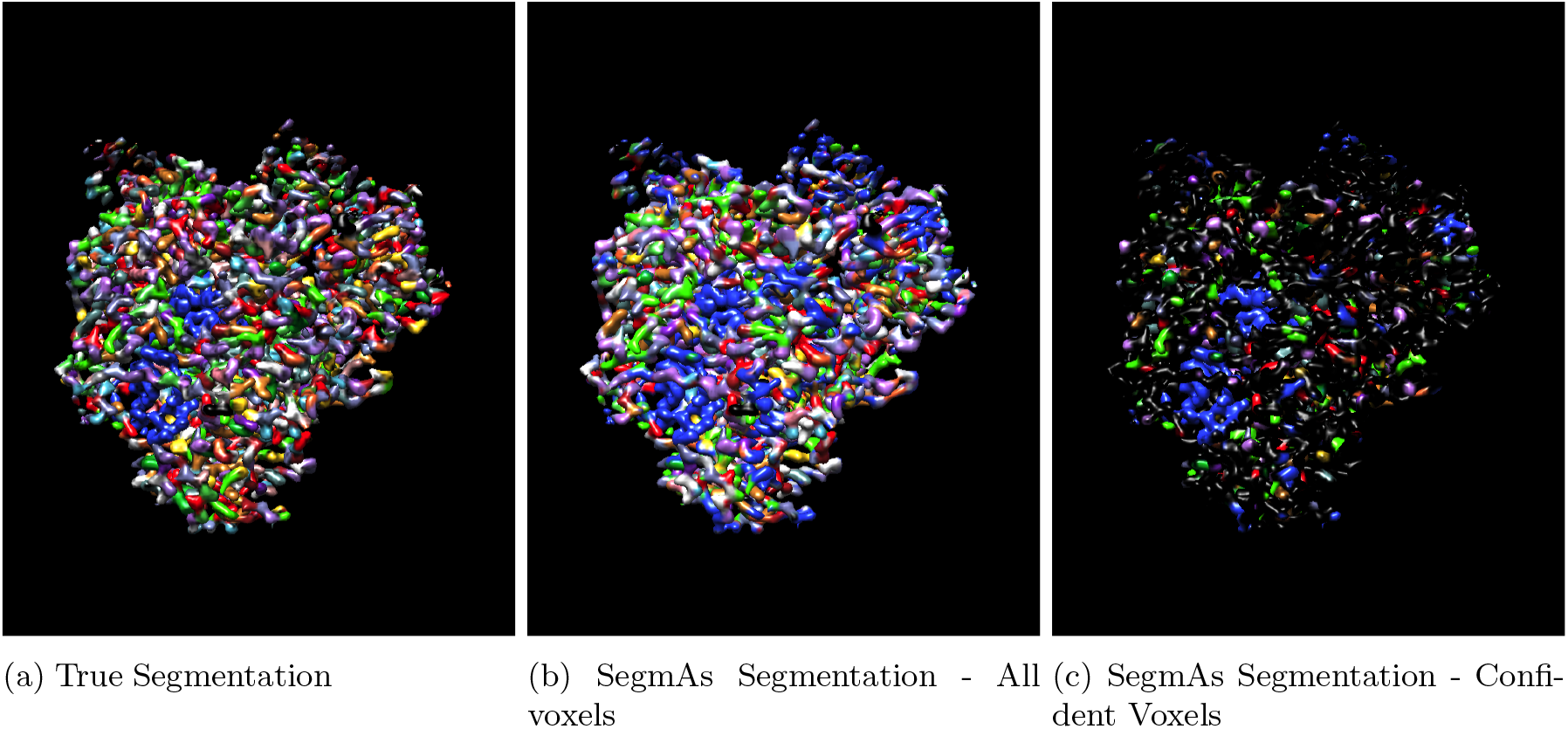
An example of unsuccessful segmentation. EMD-4512, reported map resolution - 3.2°*A*. CNF- NET removes most of the false labeled voxels. The remaining voxels belong mostly to RNA (blue), Leucine (red) and Arginine (green).

We observed that some of SegmA mislabeling cases are between amino acids which have similar properties, since they look similar on a cryo-EM map. Amino acids usually categorized to one of five groups: basic amino acids, acidic, polar uncharged, nonpolar aromatic, nonpolar aliphatic, see Fig. 14 (a). SegmAs abiliy to label correctly an amino acid group is shown at Fig. 14. As expected, SegmAs precision is considerably higher on a segmentation to groups. For example a voxel of an aliphatic amino acid can be estimated with accuracy of about 75% which can be improved to 90% if only confident voxels are considered.

**Figure 14:**
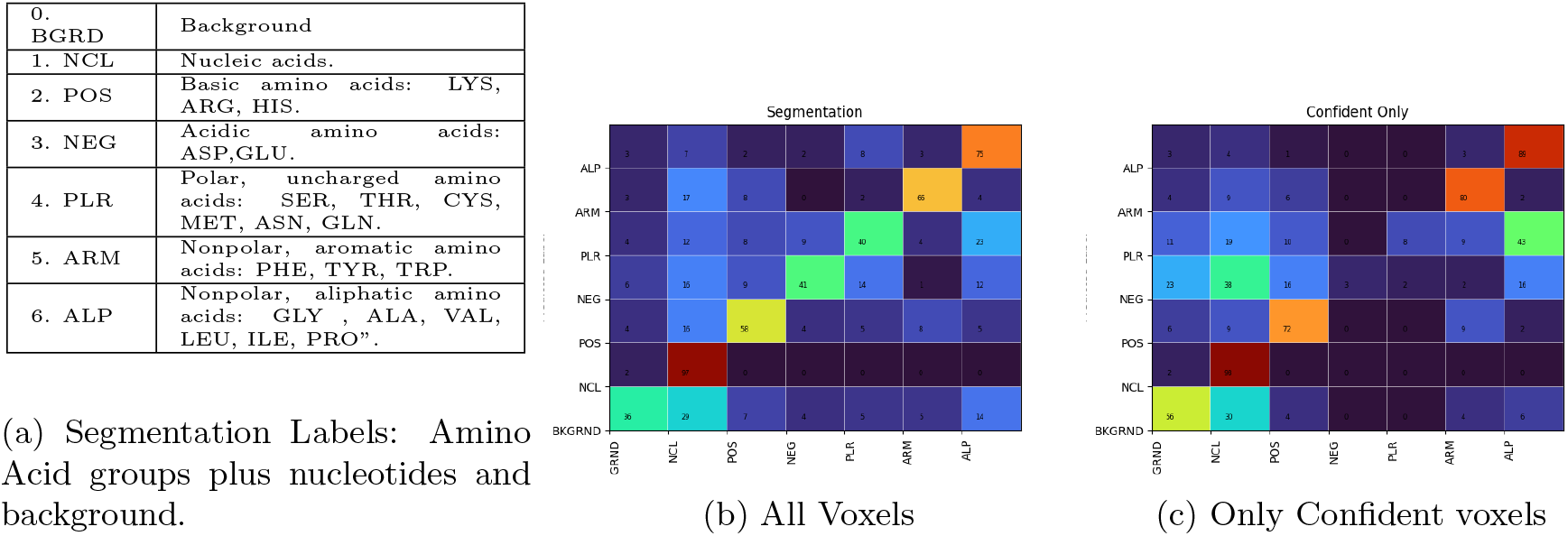
Segmentation by an amino acid groups classes.(a) Groups Definition. (b), (c) Row Normalized Confusion Matrix. A row represents a true labeling of a voxel, whereas a column represents SegmAs prediction. (a) shows the output of SEG-NET, (b) shows the results after eliminating unconfident voxels.

### 4.3 Local Resolution Effect

Although there was an anticipation for performance degradation for maps with lower resolution, one cannot see significant difference between 2.9 *−* 3.2°*A* maps and 3.3 *−* 3.5°*A* maps - Fig. 8. This is due to resolution changes within a map. In order to assess SegmAs performance as a function of resolution we estimated local resolution by Q-Score (Pintilie & Chiu (2021)). The Q-Score of an atom is a measure of how well the atom is resolved. Q-Score can be used to asses local map resolution, since it has high correlation with the resolution. Fig. 15 shows SegmAs performance as a function of the estimated local resolution. While there is clear performance degradation as the resolution lowers for amino acids, for nucleotides the dependence on the resolution is minor. When considering only confident voxels, both the precision and recall significantly improve, at the cost of total amount of labeled voxels. Fig. 15 shows the ratio of confident voxels. For local resolutions 2.3 *−* 2.5°*A* about half of the voxels are omitted, for resolutions lower than 3.5 *−* 5°*A* more than 60% are omitted.

**Figure 15:**
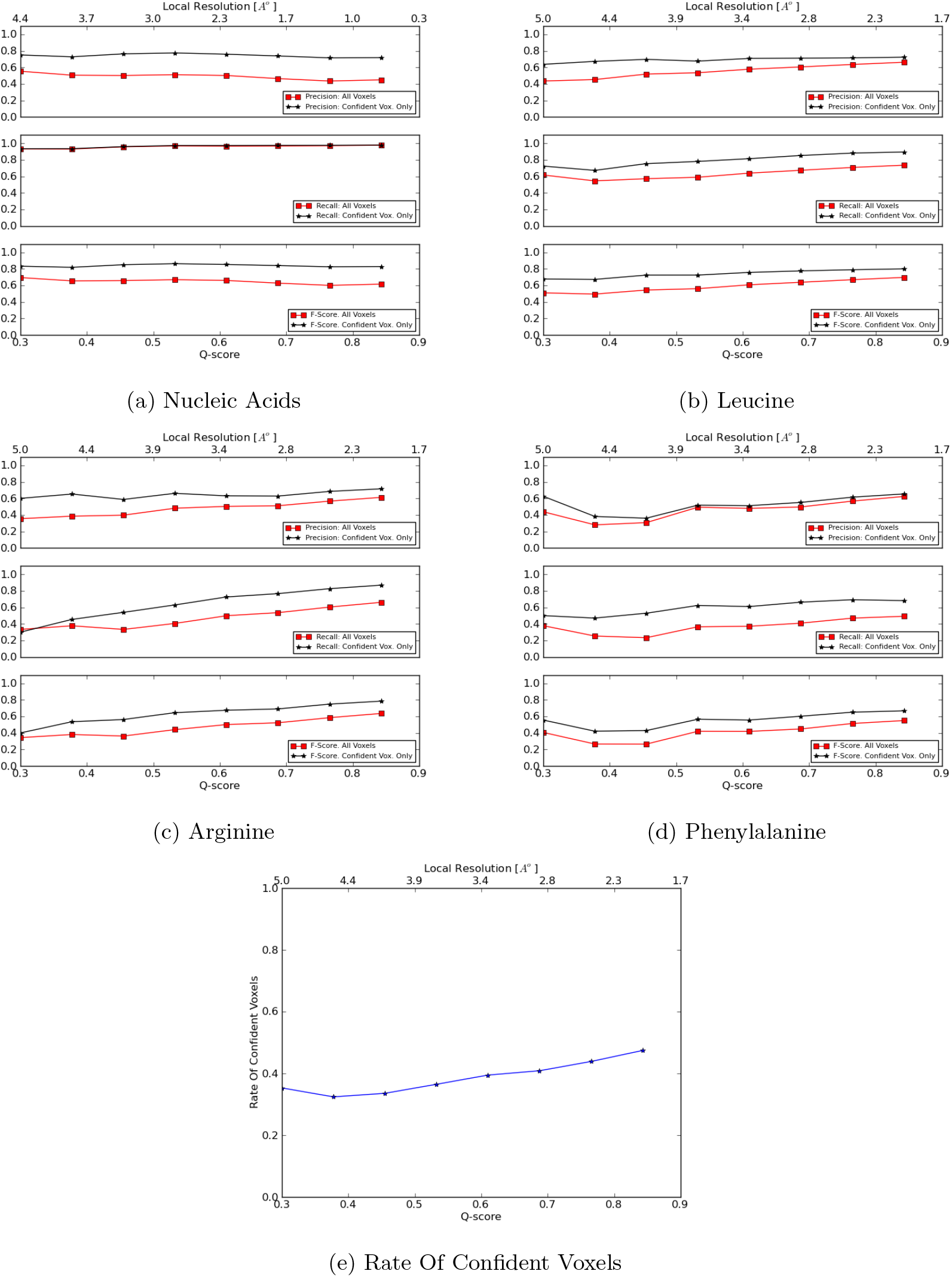
SegmAs performance as function of local resolution for selected amino acids. (a)-(d) Precision, Recall and F-Score for Nucleotides and LEU, ARG and PHE. Local resolution estimation according to Pintilie & Chiu (2021) 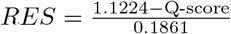 for amino acids and 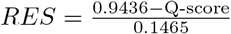 for nucleic acids.

### 4.4 Confidence Estimation

SegmaA’s performance varies for different regions of a map and different values of a map density. We also noticed from visual inspection, that CNF-NET marks as unconfident voxels which lie in a region between amino acids. Some insight on SEG-NET and CNF-NET behavior can be acquired by partitioning a map to four different regions:

- (I)- Within amino (nucleic) acids: voxels in a proximity of an atom center of given amino acids.
- (II)- AA boundary: voxels which lie in the boundary between different amino (and nucleic) acids.
- (III)- AB boundary: voxels in a boundary between an atom and the background.

A voxel in a region is assigned to one of four categories: true or false labeled by SEG-NET and labeled as confident/unconfident by CNF-NET. Fig.16 shows the ratio of voxels in each category for all the above four regions. As anticipated, the region between amino acids is characterized by low SEG-NET performance and a large amount of unconfident voxels. Since a cryo-EM map is actually an average of a large number of flexible molecules, the residue periphery often belongs to more than one amino acid or nucleotide. Moreover, voxel labeling in a periphery region is sensitive to an error in atoms positions: a small error in an atom position will not affect voxels “within amino acid”, but can cause label change for voxel between amino acids. CNF-NET successfully detects those voxels: almost 80% of voxels between amino acids were marked

**Figure 16:**
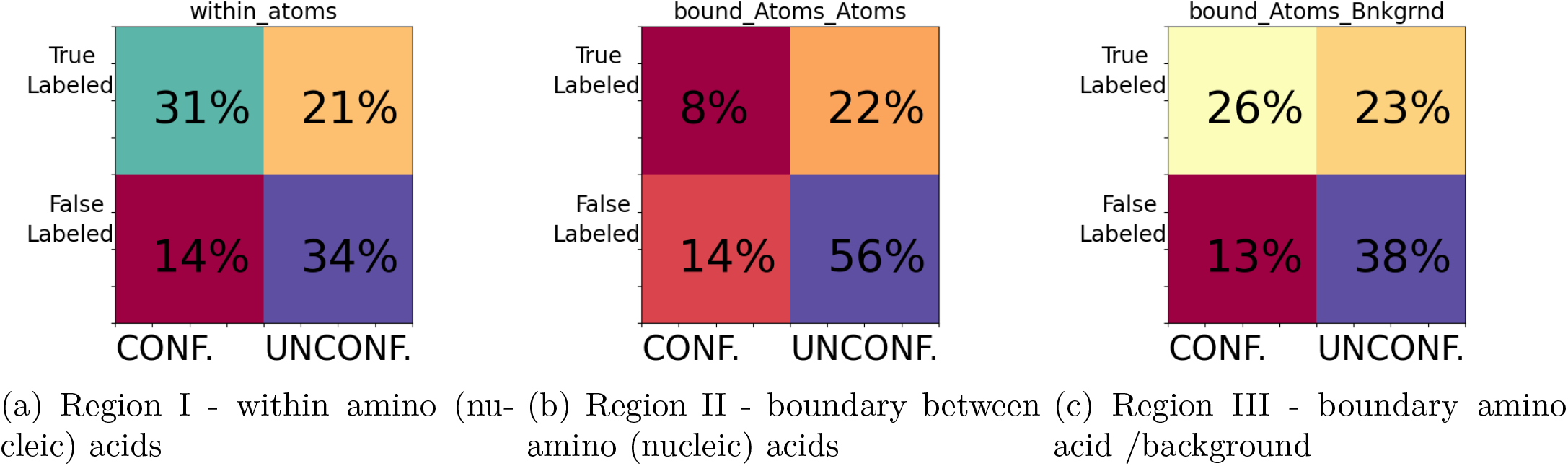
SEG-NET and CNF-NET performance for voxels in different regions.

## 5 Conclusions

SegmA is a segmentation approach to obtain structural insight into a cryo-EM map. SegmA is designed as a cascade of three convolutional networks. A fully convolutional U-net architecture enables rapid (within seconds) processing of a cryo-EM 3D map. Considerable improvement in the segmentation precision was achieved by using a group convolutional network which is the heart of the algorithm. This type of CNN has the rotational equivariance property, which leverages the native rotation invariance of a molecule. An instance segmentation task has natural performance limits when applied to a cryo EM map. Since some of the maps in the EMDB database contain regions with improper fit, those regions cannot be in a training dataset. This is done by an iterative data gathering algorithm, which created a training dataset for best segmentation performance at 3.2°*A* resolution. Some amino acids types cannot be revealed at a given resolution. Moreover, an experimental cryo-EM map has regions of degraded resolution. SegmA detects those regions and marks them as “unconfident”, which results in a considerably better performance when only confident regions are considered. SegmA estimates a residue type at the voxel level, presenting the user with a coloured 3D image. In such an image a residue pose and boundary are seen in addition to the residue position and type. In addition, SegmA has the ability to distinguish between protein residues and nucleotides.

## Acknowledgements

This research was supported by Len Blavatnik and the Blavatnik Family Foundation and by the Zimin Institute for Engineering Solutions Advancing Better Lives. Mark Rozanov was supported in part by a fellowship from the Edmond J. Safra Bioinformatics Center.

